# ULK1 and ULK2 Restrain Skeletal Myofiber Growth by Balancing Protein Synthesis and Degradation

**DOI:** 10.64898/2026.09.06.749722

**Authors:** Wangkuk Son, Jordan Fuqua, Matthew P. Harris, Ryan J. Allen, Ana Kronemberger, Julia Hughes, Luis G. O. de Sousa, Leonid Zingman, Sue C. Bodine, Benjamin F. Miller, Vitor A. Lira

## Abstract

**Background:** Skeletal muscle is vital for mobility and metabolic regulation, impacting independence and overall health. Increases in skeletal muscle mass and contractile function during development, and their maintenance during adulthood and aging, rely on an intricate coordination between protein synthesis and degradation processes that remains incompletely understood. Here, we investigated a potential role for the autophagy-initiating kinases ULK1 and ULK2 in broadly modulating protein metabolism in skeletal muscle.

**Methods:** Studies were conducted in young (4-6 wk-old) and adult (7-10 mo.-old) mice with skeletal muscle-specific knockout of *Ulk1* and *Ulk2* (i.e., Ulk1/2^skmDKO^) and wild-type littermates (WT). Short-term deficiency of these proteins was achieved via electroporation of plasmids (encoding specific microRNAs targeting *Ulk1* and *Ulk2*) into muscles of 4 mo.-old wild-type mice. Protein metabolism was assessed via deuterium oxide (D_2_O) labeling, whereas anabolic signaling was investigated under insulin and leucine administration.

**Results:** Lifelong *Ulk1/2* deficiency markedly impaired autophagy flux (i.e., LC3-II accumulated with colchicine treatment only in wild-type mice, P<0.001), compromised muscle quality, as evidenced by an increase in centrally nucleated fibers (from 0.1% to 4.5% in females, and from 0.8% to 22.7% in males (P<0.001), primarily involving MyHC type 2b fibers) and impaired force of dorsiflexors and plantar flexors in males (20%, P<0.01), and plantar flexors in females (24%, P<0.01). Despite these deficits, *Ulk1/2* deficiency promoted robust muscle hypertrophy, evidenced by increased diameters of all major MyHC fiber types in the tibialis anterior and soleus muscles (i.e., by 10-15% in males, and 14-20% in females, P<0.05). Short-term deficiency (up to 4 weeks) of *Ulk1/2* in adult skeletal muscle, however, led to myofiber hypertrophy (13%, P<0.05) without impairments in force or changes in central nucleation of fibers, pointing to an initial period of muscle quality preservation. Mechanistically, *Ulk1/2* deficiency led to elevated myofibrillar protein synthesis (23% higher Ksyn, P<0.05) and decreased mitochondrial and sarcoplasmic protein degradation (16% and 14% lower Kdeg, P=0.09 and P<0.05, respectively). Further mechanistic studies revealed that hypertrophy was accompanied by enhanced mTORC1 activity independent of AKT in *Ulk1/2*-deficient muscle.

**Conclusions:** These results indicate that ULK1 and ULK2 jointly sustain autophagy and limit mTORC1-driven protein synthesis to govern skeletal muscle protein metabolism, with lifelong deficiency increasing muscle size at the expense of quality and function, while short-term deficiency permits hypertrophy without impairment. These findings identify ULK1/2 as a novel node coordinating protein turnover in skeletal muscle, warranting investigation as a therapeutic strategy for atrophy and weakness.

## Introduction

Skeletal muscle accounts for approximately 40% of total body mass in young adults and plays a central role not only in mobility, but also in the regulation of whole-body glucose uptake and systemic protein metabolism (1-3). Skeletal muscle fibers are large, multinucleated, primarily post-mitotic cells that rely heavily on proper protein metabolism (i.e., an intricate coordination between protein synthesis and degradation processes) to preserve structural integrity and metabolic adaptability (4). In that sense, we have previously identified Unc-51-like kinases 1 and 2 (ULK1 and ULK2), the mammalian homologs of yeast Atg1 (5-7), among the few highly expressed (macro)autophagy (hereafter referred to as autophagy) genes in skeletal muscle when compared to other tissues (8); however, the degree to which ULK1 and ULK2 modulate protein metabolism in skeletal muscle remains poorly understood.

ULK1 and ULK2 share high sequence homology within their kinase domains, assemble with ATG13 and FIP200, and engage overlapping regulatory partners (9-11). Genetic evidence also suggests some functional redundancy, given that mice lacking either ULK1 or ULK2 are viable without overt autophagy defects, whereas double-null animals die perinatally (10, 12-14). Therefore, despite recent evidence supporting certain cell-specific and unique roles for ULK1 vs. ULK2, their strong expression in skeletal muscle (8, 15-17), along with the aforementioned structural, biochemical, and genetic observations, motivate a rigorous examination of their joint role in maintaining muscle proteostasis.

To carefully delineate the combined roles of ULK1 and ULK2 in skeletal-muscle proteostasis and phenotype, we studied skeletal muscle-specific Ulk1/2 double-knockout mice as well as short-term Ulk1/2 knockdown in tibialis anterior muscles via electroporation of specific microRNA (miR) plasmids targeting both *Ulk1* and *Ulk2*. In contrast to prior autophagy-deficient models, which commonly lead to muscle atrophy, we found that ULK1/2 deficiency caused myofiber hypertrophy. Mechanistically, this observation led to the discovery that ULK1/2 modulate the activity of the mammalian target of rapamycin complex 1 (mTORC1) in muscle, which, together with their direct impact on autophagy, results in a broad modulation of protein turnover pathways impacting myofiber size.

## Materials and methods

### Animals

Skeletal muscle-specific *Ulk1/2* knockout (Ulk1/2^skmDKO^) mice were generated by crossing homozygous Ulk1/2^fl/fl^ mice on a C57BL6/J background with Myogenin-Cre heterozygous mice (generous gift from Dr. Eric Olson). Offspring included Ulk1/2^skmDKO^ (Ulk1/2^fl/fl^, Myogenin-Cre^+/−^) and littermate controls (Ulk1/2^fl/fl^, Myogenin-Cre^−/−^), which were referred to as wild type (WT) mice, of varying ages and both sexes. Mice were housed under standard conditions (21°C, 12:12 h light-dark cycle) with *ad libitum* access to food and water. All experimental procedures were approved by the Institutional Animal Care and Use Committee (IACUC) of the University of Iowa.

### RNA extraction and qPCR

Total RNA (15-20 mg) from tibialis anterior (TA) and gastrocnemius (GA) muscle tissue was isolated using TRIzol reagent (Thermo Fisher Scientific, Waltham, MA, USA), followed by chloroform extraction and purified using the Direct-zol RNA Miniprep Kit (Zymo Research, Irvin, CA, USA) according to the manufacturer’s protocol. RNA quality was assessed using a Nanodrop 2000 Spectrophotometer (Thermo Fisher Scientific, Waltham, MA, USA). cDNA synthesis was performed using the High-Capacity cDNA Reverse Transcriptase Kit (Applied Biosystems, Foster City, CA, USA). qPCR was conducted with SYBR Green Master Mix (Applied Biosystems, Foster City, CA, USA) on a QuantStudio 6 Flex System.

### Colchicine administration

To inhibit autophagosome-lysosome fusion, mice received intraperitoneal injections of colchicine (0.4 mg/kg in sterile dH₂O) or vehicle at 24 and 12 h before tissue harvest (18).

### Hematoxylin and Eosin (H&E) staining

Muscle morphology and centrally nucleated fibers were assessed in H&E-stained sections. Centrally nucleated fibers were quantified as a percentage of total fibers, and specific number of fibers analyzed is provided in the figure legends. These were quantified in 7-10-month-old WT and Ulk1/2^skmDKO^ mice, and in 4-month-old WT mice 4 weeks after knockdown of *Ulk1* and *Ulk2*. Olympus BX63 microscope equipped with a DP74 camera, UPLXAPO 10x/0.40 objective, cellSens Dimension software, automated tiled acquisition of the entire muscle cross-section, with individual fields acquired at 1920 x 1920 pixels. Analysis was conducted with ImageJ/Fiji software (NIH, Bethesda, MD, USA).

### *In vivo* muscle contractile function

Muscle force production was assessed in the dorsiflexors (tibialis anterior and extensor digitorum longus) and plantar flexors (gastrocnemius, plantaris and soleus) via percutaneous stimulation of the common fibular or tibial nerve, respectively (8). Mice were positioned with their knees secured, and their feet attached to a force transducer. Stimulation (150 Hz, 300 ms) was delivered using the Aurora Scientific 1300A System, and muscle torque was analyzed using Dynamic Muscle Analysis software (version 5.321; Aurora Scientific Inc., Aurora, ON, Canada).

### Muscle sections and fluorescence microscopy

Harvested muscles were rapidly frozen by immersion in pre-cooled isopentane for ∼20 s and stored at -80 °C until sectioning. For cryosectioning, muscles were embedded in OCT using an isopentane bath to prepare blocks, then sectioned at 10 µm using a cryostat (Microm HM505E, Microm International, Walldorf, Germany). For fiber typing, sections were post-fixed in 100% acetone, permeabilized with PBS + 1% Tween 20, and blocked using M.O.M. solution (Vector Laboratories, Burlingame, CA, USA) and normal goat serum (Sigma-Aldrich, St. Louis, MO, USA). Primary antibodies included MyHC Type-I (BA-F8, 1:250), Type-IIa (SC-71s, 1:250), Type-IIb (BF-F3s, 1:250) (Developmental Studies Hybridoma Bank, University of Iowa, Iowa City, IA, USA), and Laminin (L9393, 1:500) (Sigma-Aldrich, St. Louis, MO, USA). Sections were then incubated with appropriate fluorophore-conjugated secondary antibodies for 1 h, followed by mounting with ProLong Gold Antifade Mountant without DAPI (P36930; Thermo Fisher Scientific, Waltham, MA, USA). For identification of centrally nucleated fibers per myofiber type, sections were stained for laminin, MyHC Type IIa, and MyHC Type IIb, with laminin detected using Alexa Fluor 647-conjugated donkey anti-rabbit IgG (A31573; Thermo Fisher Scientific, Waltham, MA, USA). Nuclei were counterstained with DAPI using VECTASHIELD Antifade Mounting Medium with DAPI (H-1200; Vector Laboratories, Burlingame, CA, USA). Imaging was performed using a Zeiss LSM710 Confocal Microscope. Muscle fiber diameter was analyzed using minimal Feret’s diameter via ImageJ/Fiji software (NIH, Bethesda, MD, USA) (19).

### Muscle electroporations

Plasmids encoding Ulk1 and Ulk2 micro RNAs (miRNAs), and a control miRNA encoding a nontargeting pre-miR hairpin sequence, were cloned into pcDNA6.2-GW/EmGFP-miR (Thermo Fisher Scientific, Waltham, MA, USA) as previously described (8). Essentially, TA muscles were injected with a total of 40 µg DNA plasmids (20 µg plasmid encoding *Ulk1* miRNA + 20 µg plasmid encoding either *Ulk2* miRNA) or control miRNA, followed by electroporation (175 V/cm, 10 pulses, 20 ms, 480-ms intervals) using an ECM-830 electroporator (Harvard Apparatus, Holliston, MA, USA). Muscles were harvested 1-4 weeks post-electroporation.

### Deuterium Oxide (D_2_O) labeling and assessment of protein synthesis and degradation rates

Protein synthesis and degradation rates were determined using D₂O isotope labeling, as previously described (20). Mice received an intraperitoneal bolus of 99% D₂O (0.9% NaCl, w/v), equivalent to 5% of the body water pool (considered to be at 60% of body weight), followed by ad libitum access to 8% D₂O-enriched drinking water for two weeks before sacrifice. Powdered muscle tissue (40 mg) was homogenized (1:10) in isolation buffer (100 mM KCl, 40 mM Tris-HCl, 10 mM Tris Base, five mM MgCl₂, one mM EDTA, one mM ATP, pH 7.5) containing protease and phosphatase inhibitors (Roche Diagnostics, Basel, Switzerland; Sigma-Aldrich, St. Louis, MO, USA). Hydrolyzed proteins were analyzed on an Agilent 7890A GC-MS (Agilent Technologies, Santa Clara, CA, USA). Serum D₂O enrichment was measured via liquid water isotope analysis (Los Gatos Research, Mountain, CA, USA) and was used for the precursor enrichment using mass isotopomer distribution analysis (MIDA). The protein synthesis and degradation rates were calculated using previously published methods (21).

### *In vivo* assessment of AKT- and mTORC1-dependent signaling

AKT- and mTORC1-dependent signaling was assessed *in vivo* by inspecting the anabolic response to simulated feeding. Briefly, male WT and Ulk1/2^skmDKO^ mice (6 weeks old) were fasted overnight (11-12 h), and then anesthetized with isoflurane (∼3%) in breathing air through the entire procedure. First, one TA was carefully harvested (i.e., with negligible blood loss), representing the fasting condition. Subsequently, insulin and leucine were administered intraperitoneally via a single injection (saline solution containing 3U/kg insulin and 200 mg/kg leucine). The contralateral TA was then collected 10 minutes afterwards, as previously described (22), representing the fed condition. Both muscles were flash-frozen in liquid nitrogen for subsequent analyses.

### Rapamycin administration

To inhibit mTORC1 signaling, 12-week-old mice were treated with rapamycin after TA muscle electroporations to cause miRNA-mediated *Ulk1* and *Ulk2* knockdown. Plasmids encoding *Ulk1* and *Ulk2*-targeting miRNAs were delivered into TA muscles as denoted in the Muscle electroporations section. Beginning one day after electroporation, rapamycin was administered as previously described (23). Essentially, mice received intraperitoneal injections every 24 h for six consecutive days at a dose of 0.6 mg/kg (0.6 µg/g) body weight. Rapamycin was prepared from a 5 mg/mL stock solution and diluted immediately before injection, using saline as the vehicle. Vehicle-treated mice received the corresponding solution without rapamycin (i.e., saline only). Injection volumes were calculated based on individual body weight. TA muscles were harvested 7 days after electroporation, 24 h after the final injection.

### Immunoblot analysis

Muscle samples were snap-frozen in liquid nitrogen, pulverized, and homogenized in ice-cold protein lysis buffer containing 50 mM Tris-HCl (pH 6.8), 1% SDS, 10% glycerol, 20 mM DTT, 127 mM 2-mercaptoethanol, 0.01% bromophenol blue, supplemented with protease and phosphatase inhibitors (Roche Diagnostics, Basal, Switzerland; Sigma-Aldrich, St. Louis, MO, USA). Lysates were heated at 95°C for 5 minutes, centrifuged at 15,000 × *g* for 5 minutes, and the supernatants were collected and stored at −80°C. Protein concentration was determined using the RC DC assay (Bio-Rad Laboratories, Hercules, CA, USA). Equal amounts of protein (30 μg per lane) were resolved using SDS-PAGE, transferred to PVDF, and blocked in 5% non-fat milk or BSA in TBS-T for one hour. Specific information on antibodies used is provided in Supplemental Methods.

### Proteolytic activities of the proteasome and lysosome

Proteasomal (20S and 26S) and lysosomal (cathepsin L and B) enzymatic activities were measured as described previously (8, 24, 25), with minor modifications. Specific information is provided in Supplemental Methods.

### Statistical analysis

All data were analyzed using GraphPad Prism v.10. Results are presented as means ± SEM, with sample sizes specified in the figure legends. Normality was assessed using the Shapiro-Wilk test. Comparisons between two independent groups were performed using unpaired two-tailed *t*-tests, whereas paired two-tailed *t*-tests were used for matched or contralateral-limb comparisons. Two-way ANOVA followed by the Bonferroni multiple-comparisons test was used when appropriate. Pearson correlation analysis was used to assess associations between continuous variables. The statistical test used for each dataset is specified in the corresponding figure legend. Statistical significance was set at *P* < 0.05.

## Results

### ULK1/2 deficiency leads to myofiber hypertrophy

To investigate the combined functions of ULK1 and ULK2 in skeletal muscle, we first generated skeletal muscle-specific Ulk1/2 double knockout (Ulk1/2^skmDKO^) mice, which presented decreases of 85-90% in *Ulk1* and *Ulk2* mRNA in both sexes (**Fig. 1A, B**). Body mass was ∼9% lower in adult male Ulk1/2^skmDKO^ mice, but not in females (**Fig.1C**). This was accompanied by significantly smaller perigonadal (gWAT), subcutaneous (sWAT), and brown adipose tissue (BAT) depots in Ulk1/2^skmDKO^ male mice, with a similar trend observed in females (**Suppl. Fig.1A, B**). Interestingly, Ulk1/2^skmDKO^ mice presented higher hindlimb muscle masses (15-26% in males and 10-31% in females), which included the gastrocnemius (GA), plantaris (PL), soleus (SOL), tibialis anterior (TA), and extensor digitorum longus (EDL) (**Fig.1D, E**).

**Figure 1.**
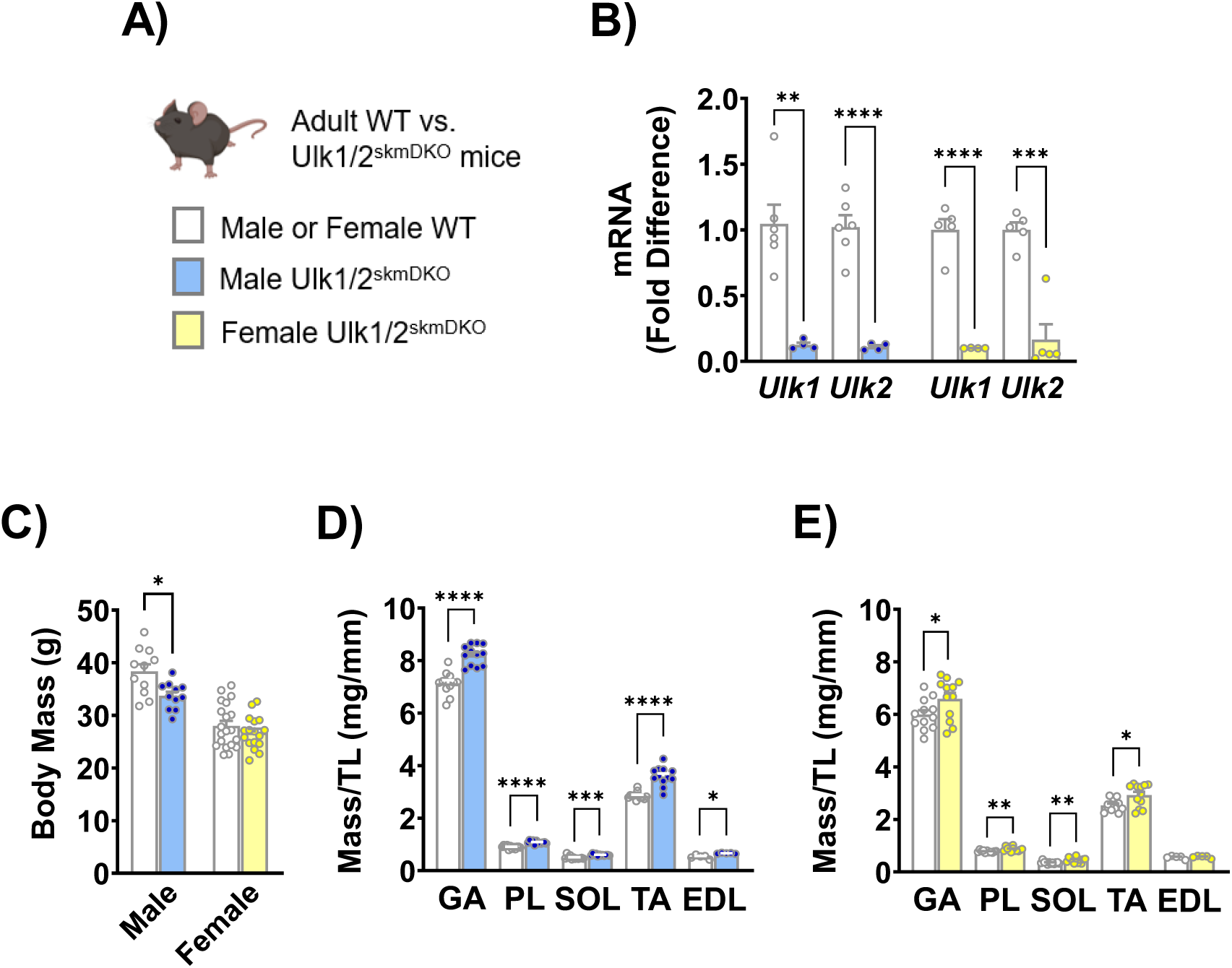
Loss of *Ulk1/2* increases muscle mass. A) Schematic representation of the experimental groups with male and female mice color-coded (Created with BioRender.com). Wild-type (WT) mice were Ulk1/2^fl/fl^, and skeletal muscle-specific Ulk1/2 double knockout (skmDKO) mice were generated by crossing ULK1/2^fl/fl^ mice with Myo-Cre^+/-^. Mature adult mice (7-10 months old) were used. B) Relative *Ulk1* and *Ulk2* mRNA expression in gastrocnemius (GA) muscle (fold difference vs. WT, n = 4-6). C) Body mass (n = 11-20). D-E) Gastrocnemius (GA), plantaris (PL), soleus (SOL), tibialis anterior (TA), and extensor digitorum longus (EDL) muscle masses normalized to tibia length (TL) in male (D, n = 9-12) and female (E, n = 5-19) mice. Data represent means ± SEM. Statistical comparisons were performed using unpaired two-tailed *t*-tests. \**P* < 0.05, \*\**P* < 0.01, \*\*\**P* < 0.001, \*\*\*\**P* < 0.0001.

Since all these muscles contain different myofiber types, we next examined whether the large muscle masses seen in Ulk1/2^skmDKO^ mice resulted from changes in myofiber size and/or distribution of specific MyHC isoforms. To address this question, we examined the TA muscle, containing primarily MyHC II fibers, and the SOL muscle, containing both MyHC II and MyHC I fibers. Ulk1/2^skmDKO^ mice had significantly larger myofiber diameters in the TA and SOL (i.e., by 10-15% in males, and by 14-20% in females, respectively). This increase affected all myofiber types and muscles, except for MyHC IIx in females, which only showed a trend for an increased diameter (**Fig. 2A-D**). The distribution of myofiber types was also altered by loss of *Ulk1/2* in muscle. A common feature was a 45-59% decrease in the proportion of MyHC IIa myofibers in the TA of both males and females. This was paralleled by an increased proportion of MyHC IIx myofibers in males (28%) and females (29%). In the more oxidative SOL muscle, a trend was seen only for a decrease in the proportion of MyHC I (8%) in Ulk1/2^skmDKO^ males (**Fig. 2A, C**). Analysis of MyHC transcripts revealed lower levels of *Myhc I* and *IIa* mRNA in the TA muscles of male and female Ulk1/2^skmDKO^ mice (**Suppl. Fig 2**). Since both the size and distribution of different MyHC myofibers within a given skeletal muscle may affect contractile function, we next assessed *in vivo* torque of dorsiflexor and plantar flexor muscles (**Fig. 2E, F**). In male mice, dorsiflexor absolute torque was unchanged, but relative torque (i.e., normalized to TA mass) was significantly lower by 20% in Ulk1/2^skmDKO^ compared to WT (P<0.01). In female mice, neither absolute nor relative dorsiflexor torque differed between genotypes. In the plantar flexors, however, both absolute and relative torques were significantly lower in Ulk1/2^skmDKO^ mice irrespective of sex (20% in males and 24% in females, P<0.01). We then examined whether myofiber structure was altered in Ulk1/2^skmDKO^ mice and observed an increased proportion of centrally nucleated fibers in the TA, but not in the SOL muscle, of both male and female Ulk1/2^skmDKO^ mice. The fact that the SOL muscle was protected suggested that MyHC IIb myofibers, which are not present in the SOL, were likely more affected. Further examination of TA muscles indeed confirmed a higher susceptibility for MyHC IIb myofibers (and lower susceptibility for MyHC IIa myofibers) to display central nuclei, particularly in females (**Fig. 3A-F** and **Suppl. Fig. 3**).

**Figure 2.**
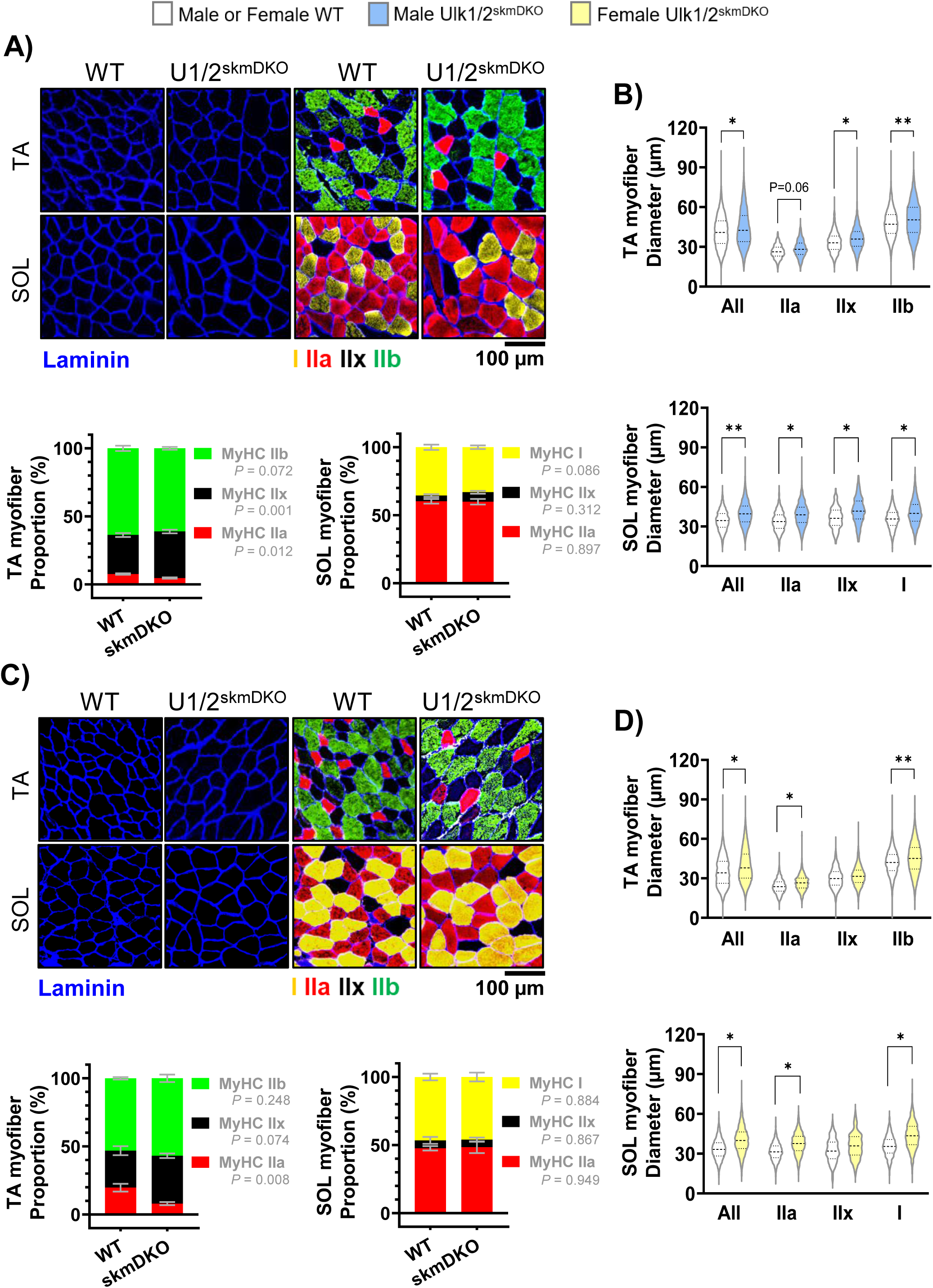
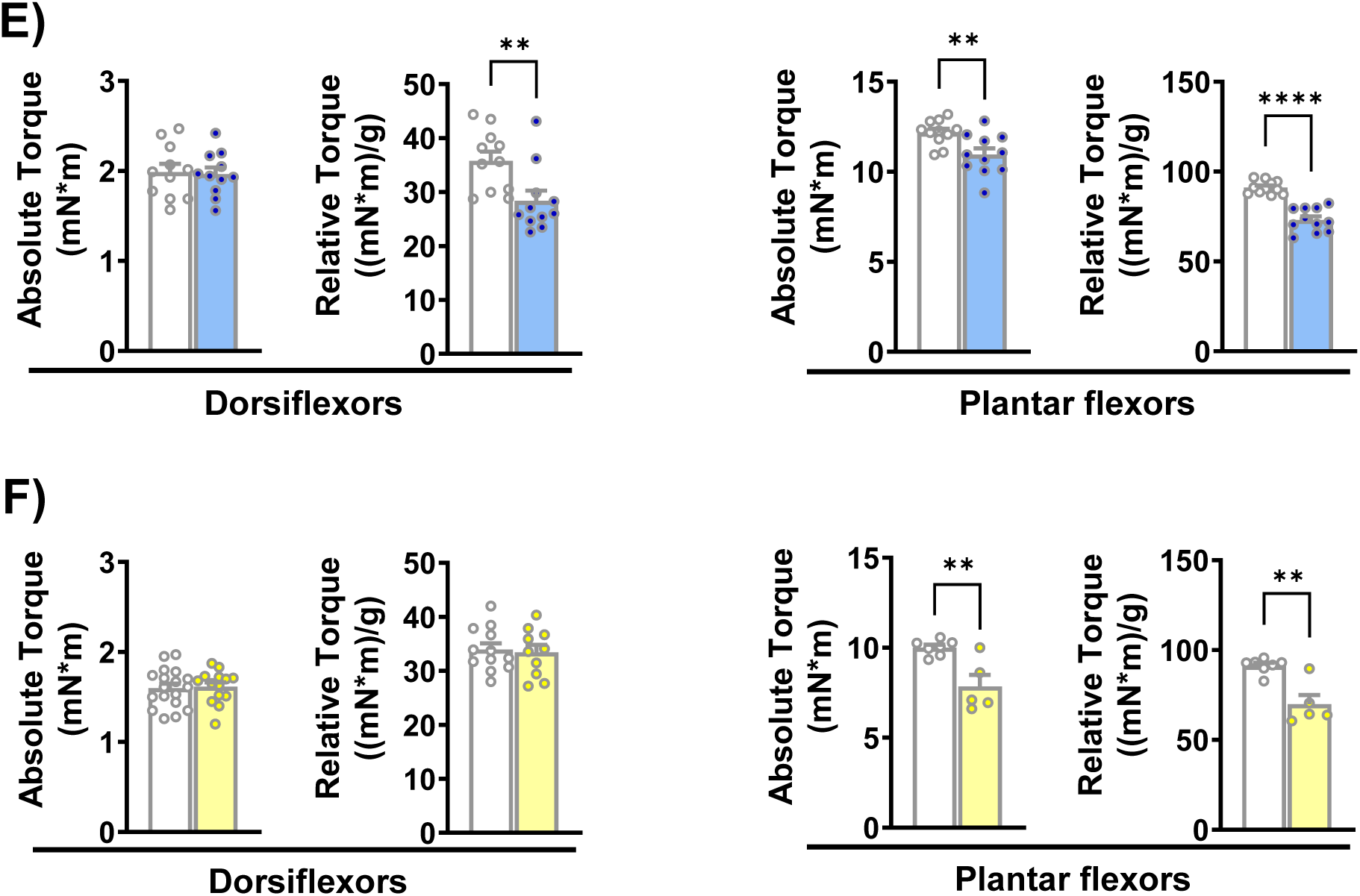
Loss of *Ulk1/2* causes broad myofiber hypertrophy without improving muscle force. A) (*Top Left*) Representative immunofluorescence images of tibialis anterior (TA) and soleus (SOL) muscles from male WT and skmDKO mice denoting laminin and MyHC isoforms (Type I, IIa, IIx, IIb). (*Bottom Left*) Fiber type proportions in TA and SOL muscles from male mice, expressed as a percentage of total fibers. Each stacked bar represents the mean ± SEM percentage of total myofibers for each myofiber type. P values shown reflect myofiber-specific comparisons between WT and skmDKO. (B) Violin plots (displaying 1^st^ quartile, median, and 3^rd^ quartile) of myofiber diameter distributions in TA (top) and SOL (bottom) muscles from male mice. For TA, WT (n = 5, total 8,687 fibers) and skmDKO (n = 5, total 10,592); for SOL, WT (n = 6, total 4,967 fibers) and skmDKO (n = 8, total 5,151 fibers). C-D) The same analyses shown in A-B were performed in female WT and skmDKO mice. For TA, WT (n = 4, total 8,216 fibers) and skmDKO (n = 4, total 9,090 fibers); for SOL, WT (n = 5, total 3,836 fibers) and skmDKO (n = 5, total 3,259 fibers). E) Absolute and relative maximal isometric torque of the ankle dorsiflexors and plantar flexors, respectively, in male mice (n = 11). F) Absolute and relative maximal isometric torque of the ankle dorsiflexors (n = 14-18) and plantar flexors, respectively, in female mice (n = 5-7). Relative torque was normalized to TA and GA muscle mass for dorsiflexors, respectively. Data represent means ± SEM. Statistical comparisons were performed using unpaired two-tailed *t*-tests. \**P* < 0.05, \*\**P* < 0.01, \*\*\*\**P* < 0.0001.

**Figure 3.**
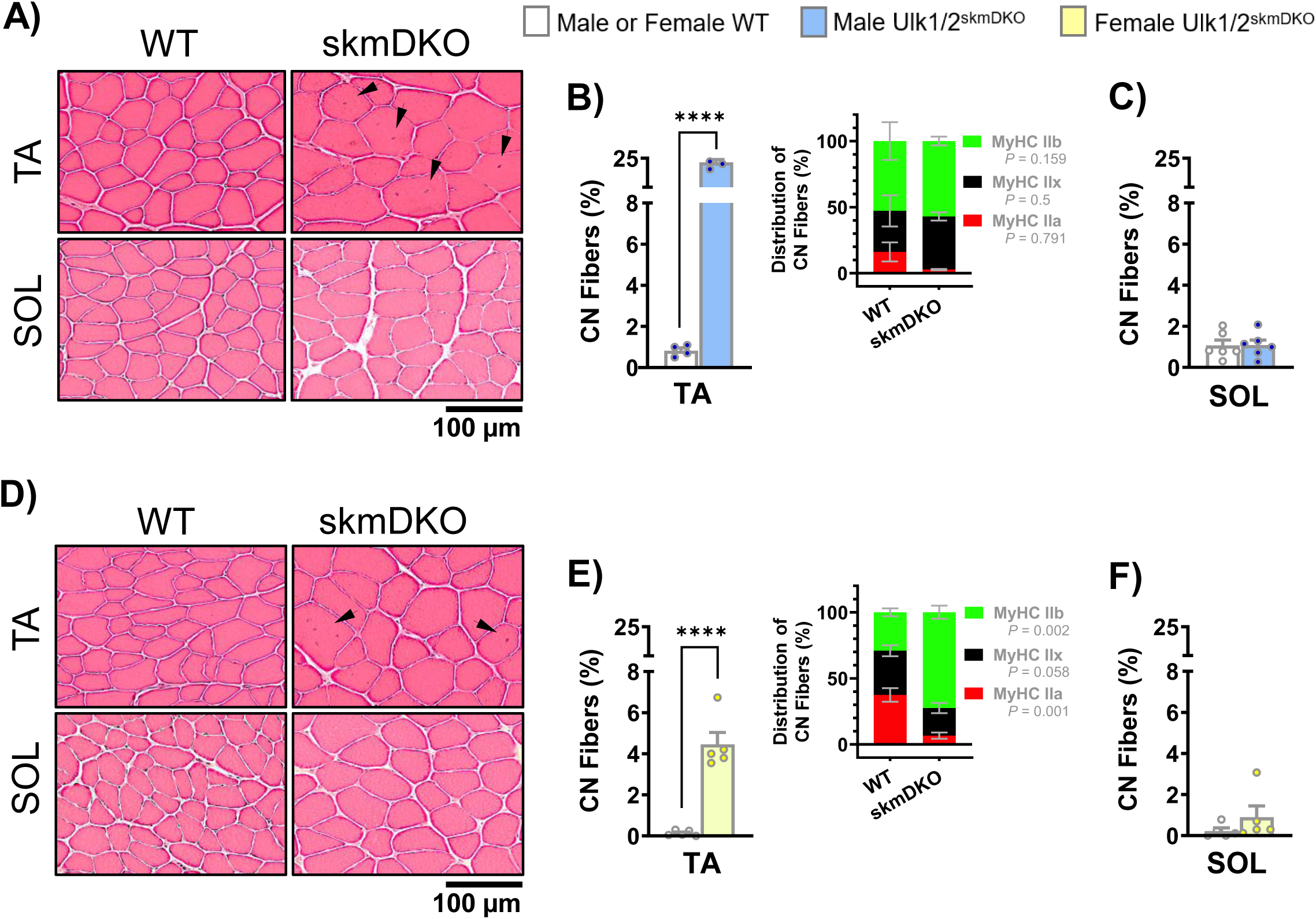
Life-long ULK1/2 deficiency increases central nucleation of MyHC IIb myofibers. A) Representative H&E-stained images showing centrally nucleated myofibers, indicated by black arrowheads, in the tibialis anterior (TA) and soleus (SOL) muscles from male WT and skmDKO mice. B-C) Quantification of centrally nucleated myofibers as a percentage of total myofibers in TA (B) and SOL (C) muscles from male mice. Myofiber type proportions of centrally nucleated myofibers in TA are shown in B. Each stacked bar represents the mean ± SEM percentage of centrally nucleated myofibers within each myofiber type. P values shown reflect myofiber-specific comparisons between WT and skmDKO. For male TA, WT (n = 4, total 7,927 fibers) and skmDKO (n = 4, total 6,980 fibers); for male SOL, WT (n = 6, total 3,989 fibers) and skmDKO (n = 6, total 3,862 fibers). D-F) The same analyses shown in A-C were performed in female WT and skmDKO mice. For female TA, WT (n = 5, total 10,382 fibers) and skmDKO (n = 5, total 9,945 fibers); for female SOL, WT (n = 5, total 3,570 fibers) and skmDKO (n = 5, total 3,793 fibers). Data represent means ± SEM. Statistical comparisons were performed using unpaired two-tailed *t*-tests. \*\*\*\**P* < 0.0001.

Next, to investigate whether the observed hypertrophy was a direct consequence of skeletal muscle ULK1/2 deficiency or a developmental compensation resulting from the perinatal deletion of *Ulk1/2* genes, we used an electroporation-based technique to transfect adult mouse TA muscle with two plasmids, which separately encoded artificial miRs specifically targeting *Ulk1* expression (miR-*Ulk1*) and *Ulk2* expression (miR-*Ulk2*). Both plasmids also encoded for EmGFP. In each mouse, the contralateral muscle received a plasmid encoding EmGFP and a nontargeting artificial miR (miR-*Control*) that served as an intrasubject negative control, as previously shown (8) (**Fig. 4A**). As expected, *Ulk1* and *Ulk2* mRNA were ∼50% decreased one week after electroporation (**Fig. 4B**). Histological evaluation of GFP-positive myofibers four weeks after electroporation revealed a significant increase by 13% in myofiber diameter of miR-*Ulk1/2* muscles (**Fig. 4C, D**). Of note, neither torque nor the percentage of centrally nucleated fibers was altered by this short-term adult deficiency of ULKs (**Fig. 4E, F**). Collectively, these findings indicate that broad myofiber hypertrophy is a direct and robust outcome of skeletal muscle ULK1/2 deficiency.

**Figure 4.**
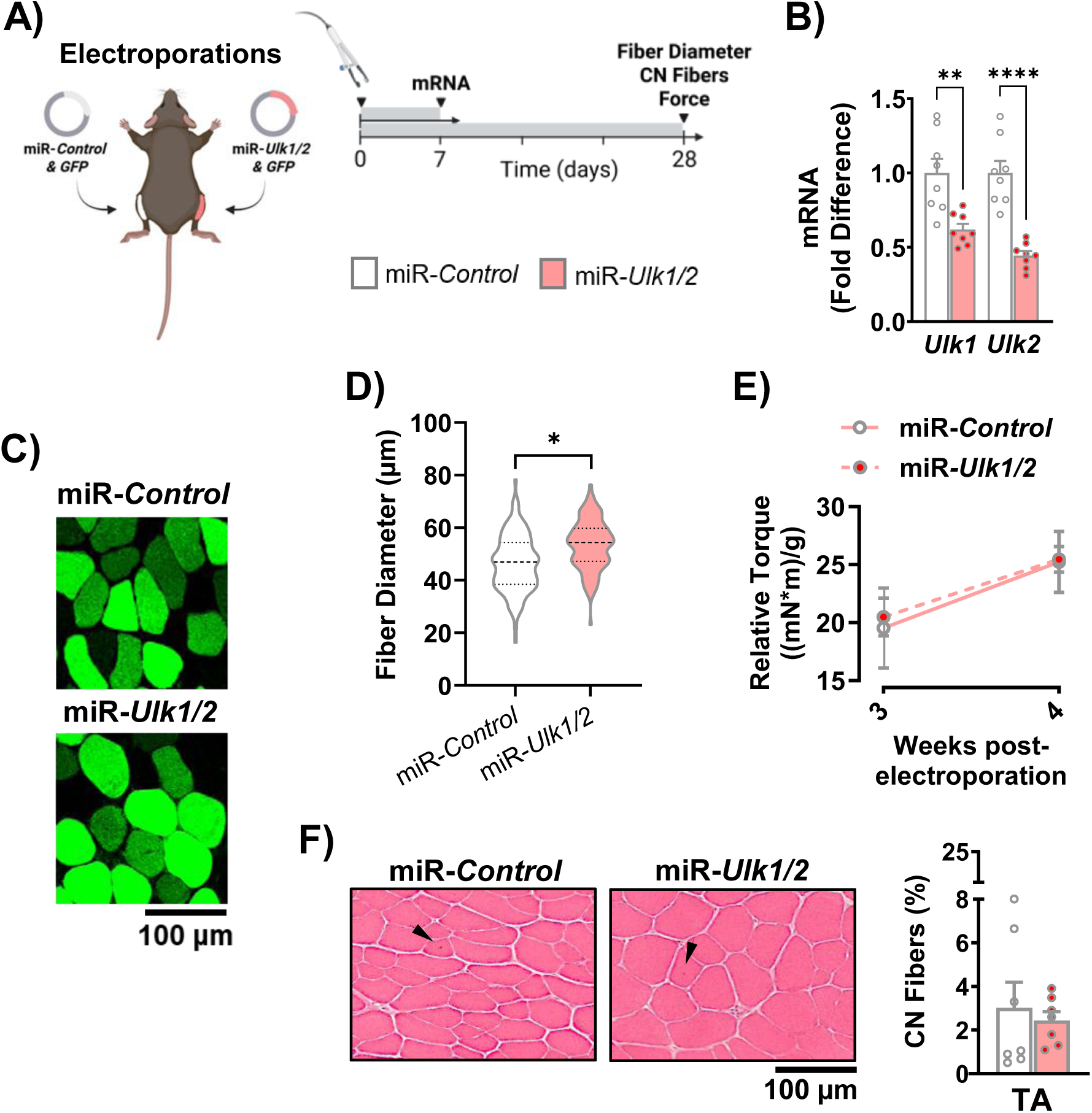
Short-term ULK1/2 deficiency increases myofiber size without impairing muscle force or increasing central nucleation. All experiments were conducted in 12-week-old male mice. A) Schematic representation of electroporations of control miR plasmid (miR-*Control*) or miR plasmids targeting *Ulk1* and *Ulk2* (miR-*Ulk1*/*2*), followed by tissue collection, functional and histological analyses in the tibialis anterior (TA) muscle (Created with BioRender.com). B) Relative *Ulk1* and *Ulk2* mRNA expression 7 days after electroporation (fold difference vs. miR-*Control*, n = 8). C) Representative images of GFP-positive fibers 28 days after electroporation. D) Violin plots (displaying 1^st^ quartile, median, and 3^rd^ quartile) of GFP-positive fiber diameter distributions 28 days after electroporation (n = 6; 120 fibers per group, with 20 fibers analyzed per muscle). E) Relative maximal isometric torque at 3 and 4 weeks post-electroporation, with torque normalized to TA muscle mass measured at the 4-week endpoint (n = 6). F) Representative H&E-stained images and quantification of centrally nucleated fibers, indicated by black arrowheads, in muscles 28 days after electroporation (n = 7; miR-*Control*, 15,695 total fibers; miR-*Ulk1/2*, 15,422 total fibers). Data represent means ± SEM. Statistical comparisons in B, D, and F were performed using paired two-tailed *t*-tests. Data in E was analyzed using two-way repeated-measures ANOVA. \**P* < 0.05, \*\**P* < 0.01, \*\*\*\**P* < 0.0001.

### Loss of ULK1/2 impairs autophagy, leading to compensatory changes in the proteasome and lysosome

Individual deficiency of ULK1 or ULK2 in skeletal muscle does not lead to broad inhibition of autophagy, which indicates that these proteins redundantly modulate basal levels of autophagy in myofibers (8, 15-17). To investigate this prospect, we examined muscle autophagy flux using colchicine, which blocks autophagosome-to-lysosome fusion, thereby leading to accumulation of the autophagosome membrane marker LC3-II. Contrary to WT mice, colchicine resulted in a subtle accumulation of LC3-II in Ulk1/2^skmDKO^ male and female mice, indicating impaired autophagy flux. Levels of ubiquitinated proteins, and to a lesser extent of p62/SQSTM1, a broad autophagy receptor, also accumulated exclusively in WT mice (**Fig. 5A, B**).

**Figure 5.**
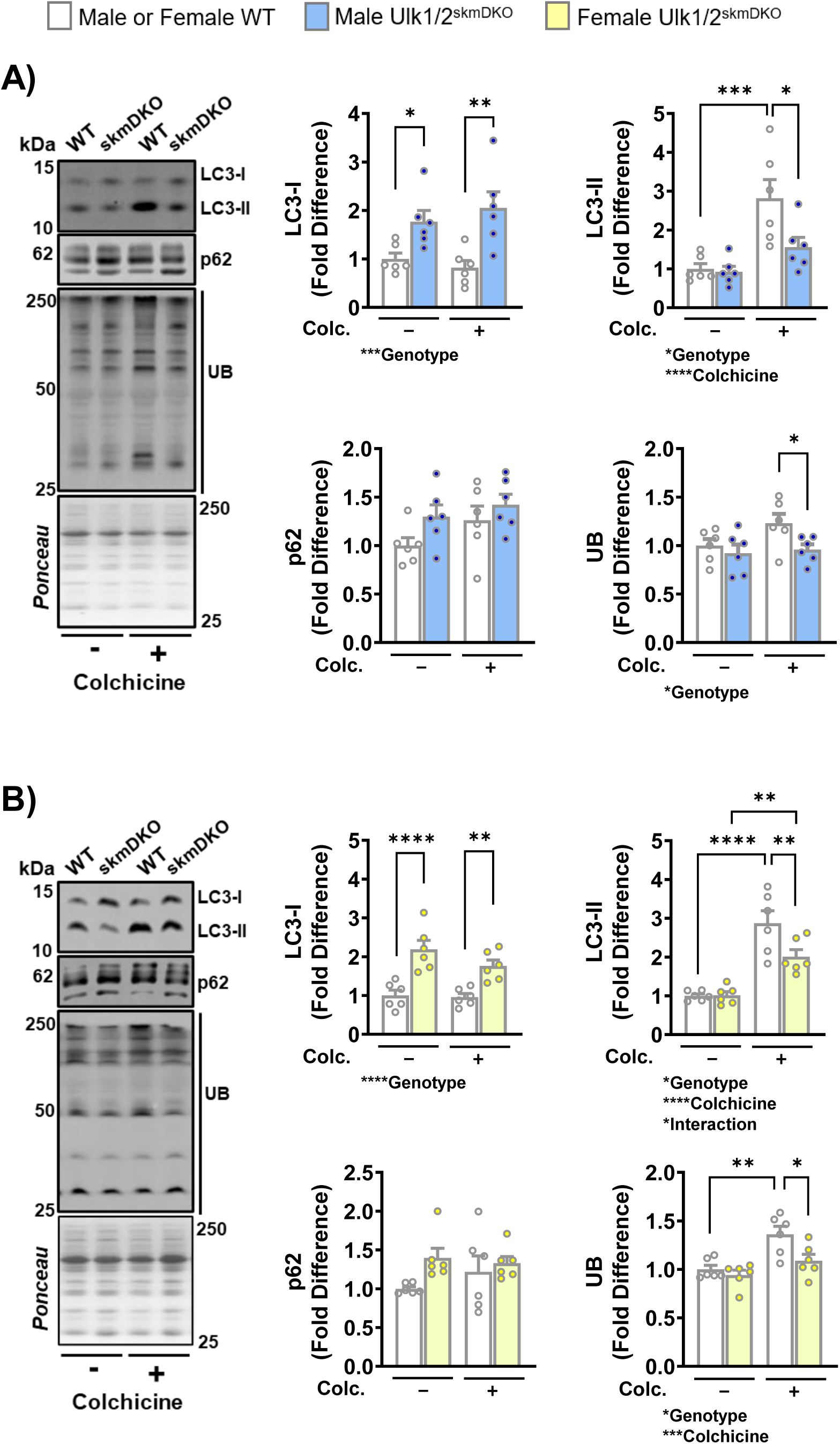
Loss of *Ulk1/2* impairs autophagy in skeletal muscle. All experiments were conducted in 12-week-old male and female mice. A) Representative immunoblots and quantification of LC3-I, LC3-II, p62, and ubiquitinated proteins (UB) in tibialis anterior (TA) muscles from male WT and skmDKO mice treated with vehicle (-) or colchicine (+) (fold difference vs. WT vehicle, n = 6). B) Same analyses shown in A were performed in female WT and skmDKO mice (fold diffenrence vs. WT vehicle, n = 6). Data represent means ± SEM. Statistical comparisons were performed using two-way ANOVA followed by Bonferroni multiple comparisons tests. \**P* < 0.05, \*\**P* < 0.01, \*\*\**P* < 0.001, \*\*\*\**P* < 0.0001.

Considering the consistent impairment of basal autophagy in muscles of male and female Ulk1/2^skmDKO^ mice, we next examined potential compensatory changes in lysosomes and proteasomes. Ulk1/2^skmDKO^ mouse muscle had higher levels of the proteasome 20S protein and the lysosomal CTSL and LAMP1 proteins, suggesting higher contents of these proteolytic machineries (**Suppl. Fig. 4A**). Accordingly, maximal activities of the proteasomal β-subunits β1, β2, and β5 were significantly higher in Ulk1/2^skmDKO^ mice vs. WT (**Suppl. Fig. 4B**). Maximal activity of CTSB, but not of CTSL, was also elevated in Ulk1/2^skmDKO^ mice (**Suppl. Fig. 4C**). Altogether, these findings indicate that muscle ULK1/2 deficiency leads to compensatory increases in content and proteolytic activity of proteasomes and lysosomes.

### ULK1/2 deficiency differentially impacts protein breakdown and synthesis across subcellular compartments

Given the marked and consistent increases in muscle mass and fiber diameter observed in adult Ulk1/2^skmDKO^ mice, irrespective of sex, we sought to define when divergence between these mice and their WT littermates appeared after birth. Although absolute muscle mass was similar between groups at 4 and 6 weeks of age, the relative increase over this period was significantly greater in Ulk1/2^skmDKO^ mice, representing a relevant window of enhanced muscle growth (**Fig. 6A**). To further explore the mechanisms underlying this hypertrophic response, we employed deuterium oxide (D₂O) isotope labeling and quantified *in vivo* protein synthesis and degradation rates (**Fig. 6B**) (21). Ulk1/2^skmDKO^ mice had a higher rate of myofibrillar protein synthesis (i.e., 23%), which was accompanied by lower degradation rates of sarcoplasmic proteins (i.e., 26%) and a trend for lower degradation rates of mitochondrial proteins (i.e., 24%) (**Fig. 6C-E**). These results indicate that the hypertrophic response in Ulk1/2^skmDKO^ mice is driven by a selective enhancement of myofibrillar protein synthesis, coupled with decreased sarcoplasmic and mitochondrial protein breakdown, collectively leading to a broad increase in protein accretion.

**Figure 6.**
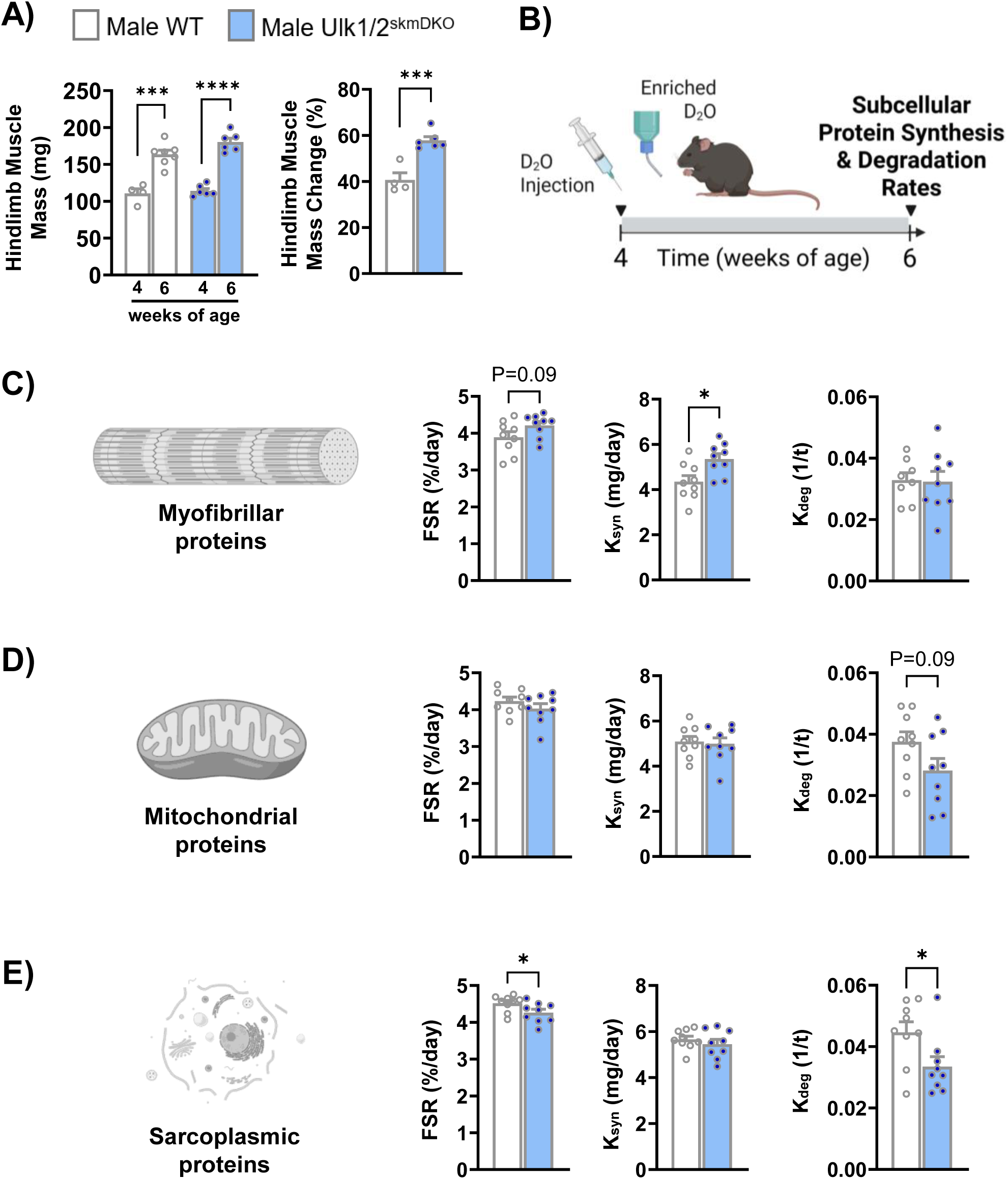
Loss of *Ulk1/2* differently impacts protein breakdown and synthesis across subcellular compartments. All experiments were performed in male mice from 4 to 6 weeks of age. A) Combined hindlimb muscle mass, including gastrocnemius, plantaris, soleus, tibialis anterior, and extensor digitorum longus, at 4 and 6 weeks of age, and the percent changes in combined muscle mass over this period (n = 4-7). B) Experimental timeline illustrating D₂O administration and tissue collection (Created with BioRender.com). C) Fractional synthetic rate (FSR), protein synthesis rate (K_syn_), and degradation rate (K_deg_) of myofibrillar proteins (n = 8-9). D) FSR, K_syn_, and K_deg_ of mitochondrial proteins. (n = 9). E) FSR, K_syn_, and K_deg_ of sarcoplasmic proteins (n = 9). Data represent means ± SEM. Statistical comparisons were performed using unpaired two-tailed *t*-tests. \**P* < 0.05, \*\*\**P* < 0.001, \*\*\*\**P* < 0.0001.

### ULK1/2 deficiency enhances protein synthesis via mTORC1 signaling

To further investigate the mechanisms underlying muscle protein accretion in Ulk1/2^skmDKO^ mice, we assessed the activity of the AKT-mTORC1 signaling pathway, which plays a central role in stimulating protein synthesis (26). Given that this pathway is tightly regulated by nutrient availability, we examined key signaling proteins under conditions of fasting and simulated feeding (i.e., following administration of insulin and leucine) in contralateral muscles of each animal (refer to methods for details). Immunoblotting showed that phosphorylation of AKT (Thr308), as well as of its downstream targets GSK-3β (Ser9) and PRAS40 (Thr246), was robustly and comparably stimulated by insulin and leucine in WT and Ulk1/2^skmDKO^ mice. However, in response to insulin and leucine, Ulk1/2^skmDKO^ mice showed increased phosphorylation of mTORC1 downstream targets (i.e., P70S6K, RPS6, and 4EBP1) compared to WT mice (**Fig. 7A**). Interestingly, a more detailed assessment of RPS6 in adult Ulk1/2 deficient muscle revealed a baseline increase in RPS6 phosphorylation at two distinct sites (S235/236 and S240/244). We then tested the extent to which this change was mediated by increased activity of mTORC1 by administering daily injections of rapamycin for 6 days after electroporations in WT mice. Our results demonstrated that phosphorylations at S235/236 and S240/244 are highly correlated across WT and Ulk1/2^skmDKO^ muscles and are comparably sensitive to rapamycin (**Fig. 7C, D**). Additionally, rapamycin completely abolished the enhanced phosphorylation at these RPS6 sites in ULK1/2 deficient muscles, further denoting the modulation of mTORC1 activity by muscle ULKs (**Fig. 7E-F**).

**Figure 7.**
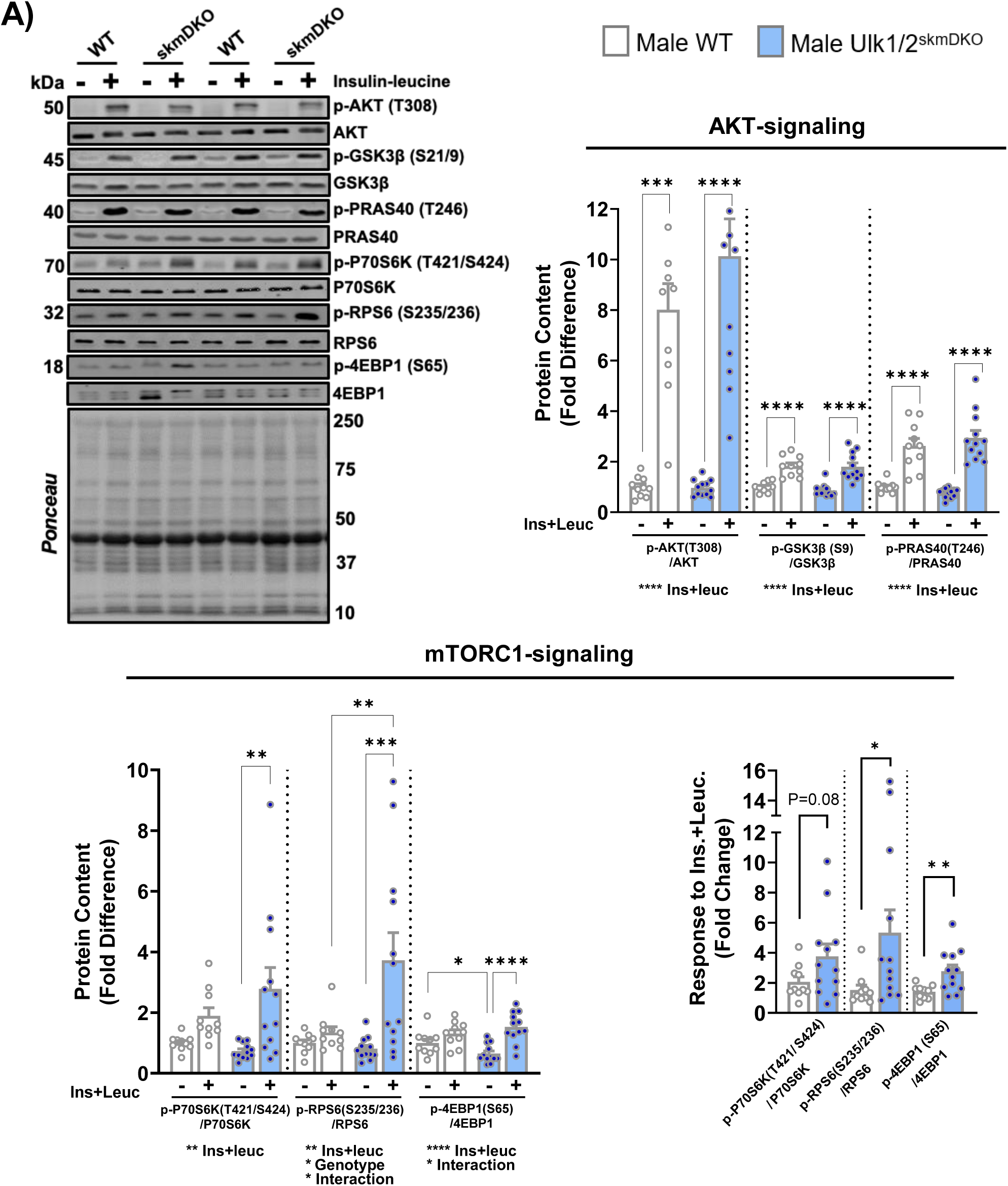
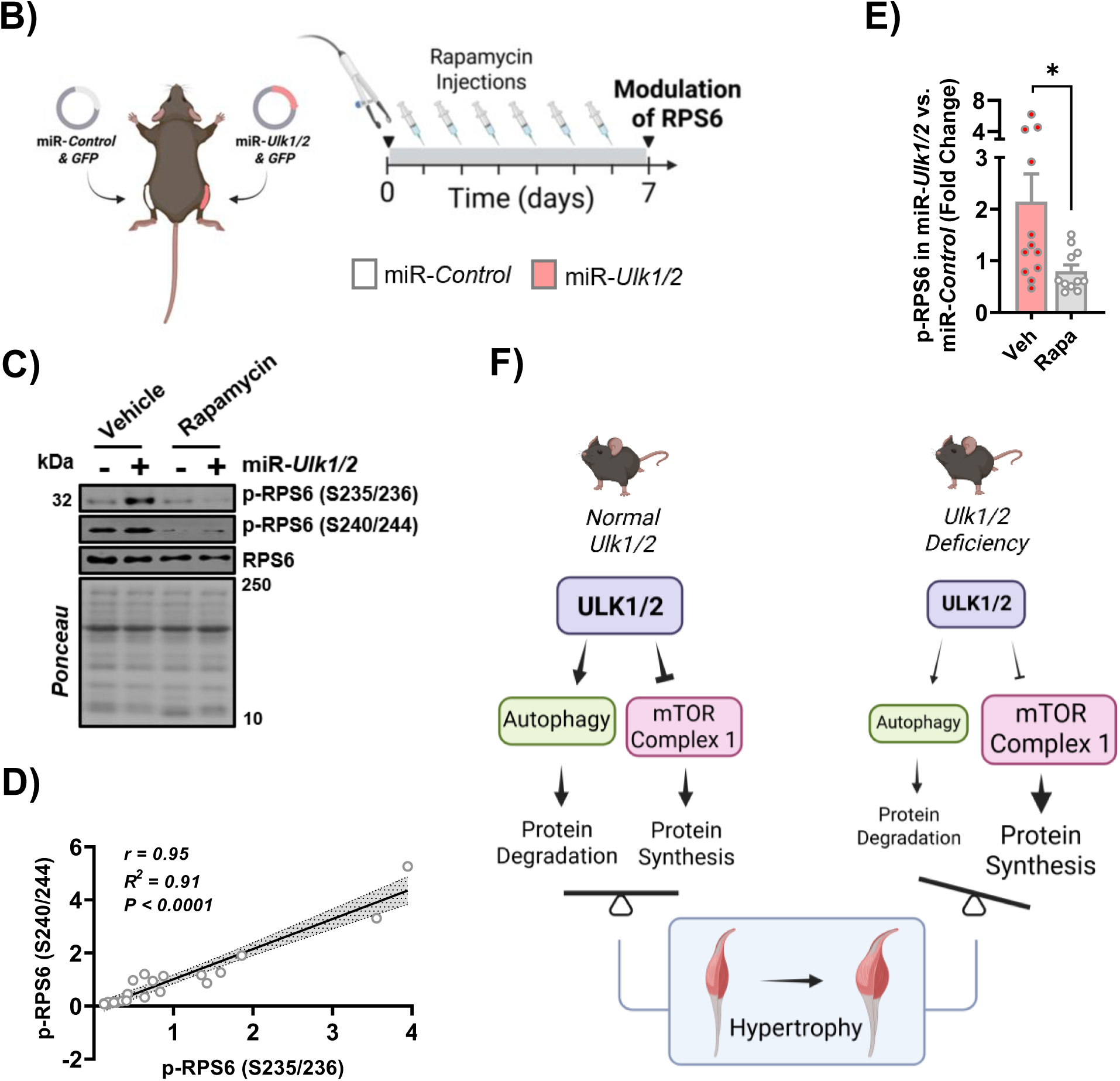
*Ulk1/2* deficiency enhances protein synthesis via mTORC1 signaling. All experiments were performed using the tibialis anterior (TA) muscle in male mice. Experiments shown in A were conducted in 6-week-old mice, whereas experiments shown in B-F were conducted in 12-week-old mice. A) (*Top Left*) Representative immunoblots of total and phosphorylated AKT, GSK3ß, PRAS40, P70S6K, RPS6, and 4EBP1 from WT and skmDKO mice under fasting (−) and insulin+leucine-stimulated (+) conditions. (*Top Right*) Quantification of phosphorylation-to-total protein ratios for AKT, GSK3ß, and PRAS40, representing AKT-signaling (fold difference vs. WT fasting condition, n = 10-12). (*Bottom Left*) Quantification of phosphorylation-to-total protein ratios for P70S6K, RPS6, and 4EBP1, representing mTORC1-signaling (fold difference vs. WT fasting condition, n = 10-12), and (*Bottom Right*) response (fold change) to insulin+leucine administration in relation to fasting condition for WT and Ulk1/2^skmDKO^ (n = 10-12). B) Schematic representation of electroporations of control miR plasmid (miR-*Control*) or miR plasmids targeting *Ulk1* and *Ulk2* (miR-*Ulk1*/*2*) in the TA muscle, followed by daily intraperitoneal injections of vehicle or rapamycin beginning 1 day after electroporation (Created with BioRender.com). Tissue collection occurred 24h after the final injection. C) Representative immunoblots of total and phosphorylated RPS6 at Ser235/236 and Ser240/244 in miR*-Control* and miR*-Ulk1/2-*electroporated muscles from vehicle- and rapamycin-treated mice. D) Pearson correlation between RPS6 phosphorylation at Ser235/236 and Ser240/244 across miR*-Control* and miR*-Ulk1/2* muscles from vehicle- and rapamycin-treated mice (n = 22). E) Combined RPS6 phosphorylation response at Ser235/236 and Ser240/244 in miR*-Ulk1/2* muscles relative to contralateral miR*-Control* muscles in vehicle- and rapamycin-treated mice (n = 11-12). F) Schematics illustrating the collective impact of muscle ULK1 and ULK2 on autophagy and mTORC1, thereby affecting protein metabolism and myofiber size (Created with BioRender.com). Data represent means ± SEM. Data in A (*Top Right* and *Bottom Left*) were analyzed using two-way ANOVA to assess the effects of genotype and insulin/leucine stimulation and their interaction, followed by Bonferroni multiple-comparisons tests. Data in A (*Bottom Right*) and E were analyzed using an unpaired two-tailed *t*-test. The association shown in D was determined using Pearson correlation. \**P* < 0.05, \*\**P* < 0.01, \*\*\**P* < 0.001, \*\*\*\**P* < 0.0001.

## Discussion

Our results demonstrate that ULK1 and ULK2 are essential regulators of protein metabolism in skeletal muscle. Interestingly, their deficiency not only impaired autophagic flux but also promoted increases in protein synthesis, leading to a robust hypertrophic response across several muscle groups. These findings reveal that, functionally, ULK1/2 differ from several other autophagy-related proteins by extending beyond their canonical autophagic role to modulate mTORC1 activity and protein synthesis, impacting myofiber size.

Combined ULK1/2 loss or deficiency caused a robust impairment of skeletal muscle autophagy flux, which was not observed with single targeted deficiencies of ULK1 or ULK2 (8, 15-17). This indicates that although each kinase may uniquely contribute to autophagy in specific settings, they operate redundantly to sustain basal autophagy flux in skeletal muscle. This autophagy impairment by ULK1/2 deficiency was accompanied by compensatory increases in the broad proteolytic capacity of the proteasome, while a more nuanced increase was seen in the lysosome (i.e., CTSB activity was increased, while CTSL was not). These findings agree with recent reports of crosstalk between autophagy and the proteasome system. Essentially, while these two major proteolytic pathways have largely non-overlapping substrate repertoires under normal conditions, inhibition of autophagy is often accompanied by compensatory upregulation of the proteasome, apparently to facilitate the clearance of cellular material that would otherwise remain unprocessed (27-29). Future investigations into the precise mechanisms underlying these adaptive responses, which specifically affect the proteasome and the lysosome, may uncover new ways to combat impairments in muscle proteostasis in certain settings, such as in metabolic diseases, aging, and inclusion body myositis (IBM).

Several previous reports have consistently linked the loss of autophagy-related genes to muscle atrophy (8, 30-33). However, despite impairing autophagy, perinatal loss of ULK1/2 in skeletal muscle led to a clear hypertrophic response encompassing different hindlimb muscles and myofiber types in both males and females (**Fig. 1D, E** and **Fig. 2A-D**). Myofiber hypertrophy was then confirmed by short-term deficiency of ULK1/2, indicating that these outcomes did not result from compensatory responses occurring during muscle development. Isotope-labeling experiments revealed that the larger myofiber diameters were driven by increased synthetic rates of myofibrillar proteins, together with decreased degradation rates of sarcoplasmic and mitochondrial proteins, collectively leading to myofibrillar hypertrophy and, possibly, some degree of sarcoplasmic hypertrophy as well (34). These observations point to a few key mechanistic prospects altering protein metabolism towards higher synthesis vs. degradation rates, thereby leading to hypertrophy in ULK1/2 deficient muscle. First, on the proteolysis side, the decreased protein degradation rates of mitochondrial and sarcoplasmic proteins, but not myofibrillar proteins, may result from basal autophagy selectively targeting mitochondrial and sarcoplasmic proteins, and/or from basal autophagy impairments affecting all the investigated protein pools, but with the increased proteasome activity being sufficient to only compensate for myofibrillar protein degradation. Second, from a protein synthesis perspective, the observed changes may reflect a selective increase in myofibrillar protein synthesis rates. Alternatively, it may reflect a more general stimulation of protein synthesis (i.e., across myofibrillar, mitochondrial and sarcoplasmic protein pools) that evades a concerted regulation coupling protein synthetic and degradation rates. This possibility is based on the fact that when compared to WT, Ulk1/2^skmDKO^ protein synthesis rates outperformed degradation rates across all protein pools (i.e., higher synthesis with similar degradation for myofibrillar proteins, and similar synthesis with lower degradation for mitochondrial and sarcoplasmic proteins). Although additional mechanistic investigations on these prospects are still required, our results reveal an important novel role for ULK1/2 proteins in jointly modulating skeletal muscle protein synthesis. These findings may have important implications for conditions of altered protein metabolism in skeletal muscle (e.g., disuse, aging, obesity, and diabetes).

The increased muscle mass resulting from perinatal deletion of skeletal muscle ULK1/2 did not translate into increased force. Our functional analyses showed that torque production by the plantar flexor muscles was significantly lower in both male and female Ulk1/2^skmDKO^ mice, while dorsiflexor function remained largely unaffected except for a reduction in relative torque (i.e., normalized to muscle mass) in males (**Fig. 2E, F**). Since muscle mass or fiber diameter are typical determinants of force generation, the unaltered or decreased force output seen in the larger muscles of Ulk1/2^skmDKO^ mice may suggest limitations in muscle quality. Accordingly, we observed an increase in the percentage of centrally nucleated fibers (i.e., undergoing damage-regeneration cycles) in the TA muscle, but not in the soleus. Additional analysis revealed that central nuclei were present primarily in MyHC IIb myofibers of Ulk1/2^skmDKO^ mice, thereby explaining why the soleus muscles, possessing mainly MyHC IIa and MyHC I fibers, were protected.

Nevertheless, it is important to note that these changes resulted from lifelong ULK1/2 deficiency and may require long periods to develop. Supporting this notion, myofiber hypertrophy was observed after four weeks of ULK1/2 deficiency in adult muscles without changes in percentage of centrally nucleated fibers or loss of force, as assessed in electroporation experiments with *Control*- and *Ulk1/2*-targeting miR plasmids (**Fig. 4**). Collectively, these observations suggest that muscle quality is preserved in all myofiber types during the early stages of ULK1/2 deficiency and myofiber hypertrophy (i.e., at least for one month in mice), and that it may remain preserved for longer in MyHC IIa and MyHC I myofibers. Although the protective mechanisms present in these myofiber types are likely multifaceted, these may be related to a higher resilience to oxidative stress, commonly resulting from impaired autophagy (31). Noticeably, these findings suggest that short-term inhibition of skeletal muscle ULK1/2 may represent a viable intervention to minimize atrophy and accelerate recovery in conditions where autophagy is stimulated and protein synthesis is impaired, such as in disuse (e.g., immobilization, hospitalization). Future studies examining this prospect are warranted.

Our findings also begin to reveal the molecular mechanisms by which ULK1/2 modulate protein synthesis in skeletal muscle. In mice with perinatal deletion of muscle *Ulk1/2*, mTORC1 activity was increased under simulated feeding conditions (i.e., administration of insulin and leucine) despite no changes in insulin-stimulated AKT activity. Of note, short-term deficiency of ULK1/2 in adult muscle also enhanced mTORC1 activity, with rapamycin treatment blocking this effect in adult myofibers. Prior studies have shown that ULK1/2 phosphorylate Raptor at inhibitory sites to limit mTORC1 activity (35-38), suggesting that the loss of this regulation may relieve inhibition and drive the heightened anabolic signaling leading to the muscle growth observed. Future studies should investigate the degree to which the hypertrophic response seen in ULK1/2-deficient muscle is mediated by Raptor-dependent activation of mTORC1 and whether it involves additional mechanisms, thereby establishing a more direct molecular link between ULK1/2, mTORC1, and control of myofiber size.

In conclusion, our study establishes ULK1 and ULK2 as indispensable kinases that jointly sustain autophagy and critically modulate anabolic signaling and muscle growth (**Fig. 7F**). While long-term deficiency of ULK1/2 throughout development promotes hypertrophy and impairs muscle quality, short-term ULK1/2 deficiency leads to hypertrophy without negatively impacting muscle quality in adult myofibers. These insights not only advance our fundamental understanding of muscle biology but also provide a new perspective on the integration between catabolic and anabolic pathways impacting skeletal muscle protein metabolism and size. These results may have therapeutic implications for a variety of conditions associated with enhanced autophagy and limited protein synthesis contributing to myofiber atrophy.

## Acknowledgments

We would like to acknowledge the University of Iowa Central Microscopy Research Facility supported by the Office of the Vice President for Research. This work was supported by the NIH (R56AG0080101), the Fraternal Order of Eagles Diabetes Research Center, and the Office of the Vice-President for Research at the University of Iowa (V.A.L.), and by the VA Career Scientist Award (IK6BX007150) (B.F.M.).

## Author contributions

V.A.L., W.S. and J.F. conceived the original idea for this study. W.S., J.F., M.P.H., R.J.A., A.K., J.H, L.G.O.S. performed experiments. S.C.B., B.F.M., and L.Z. shared expertise on specific experiments and resulting analyses. W.S., J.F., M.P.H., and V.A.L. interpreted the data. W.S. and V.A.L. wrote the manuscript. S.C.B., B.F.M., and L.Z. reviewed and revised the manuscript before submission.

## Figure Legends

**Supplementary figure 1.**
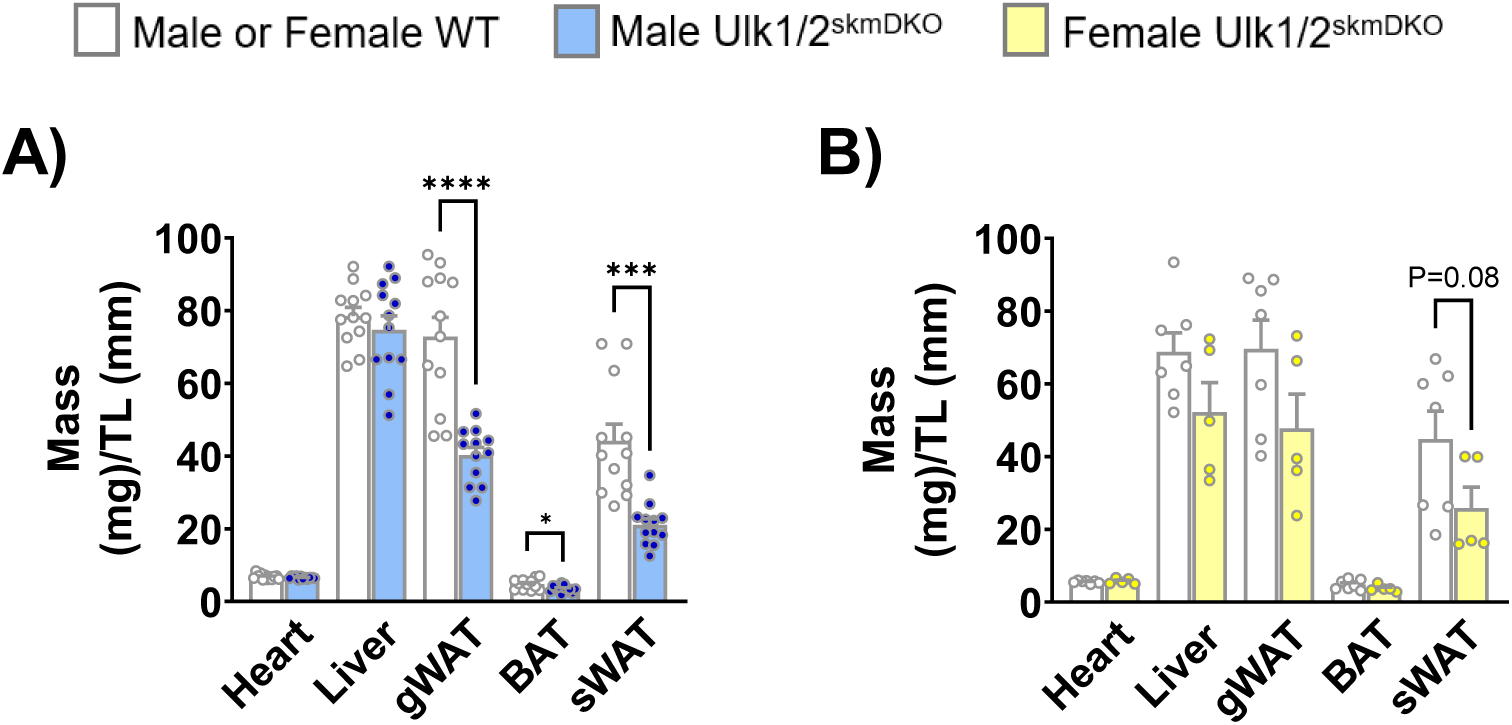
Organ and adipose tissue mass. A-B) Masses of heart, liver, gonadal white adipose tissue (gWAT), brown adipose tissue (BAT), and subcutaneous white adipose tissue (sWAT) normalized to tibia length (TL) in male (A, n = 11-12) and female (B, n = 5-7) mice. Data represent means ± SEM. Statistical comparisons were performed using unpaired two-tailed *t*-tests. \*\*\**P* < 0.001, \*\*\*\**P* < 0.0001.

**Supplementary figure 2.**
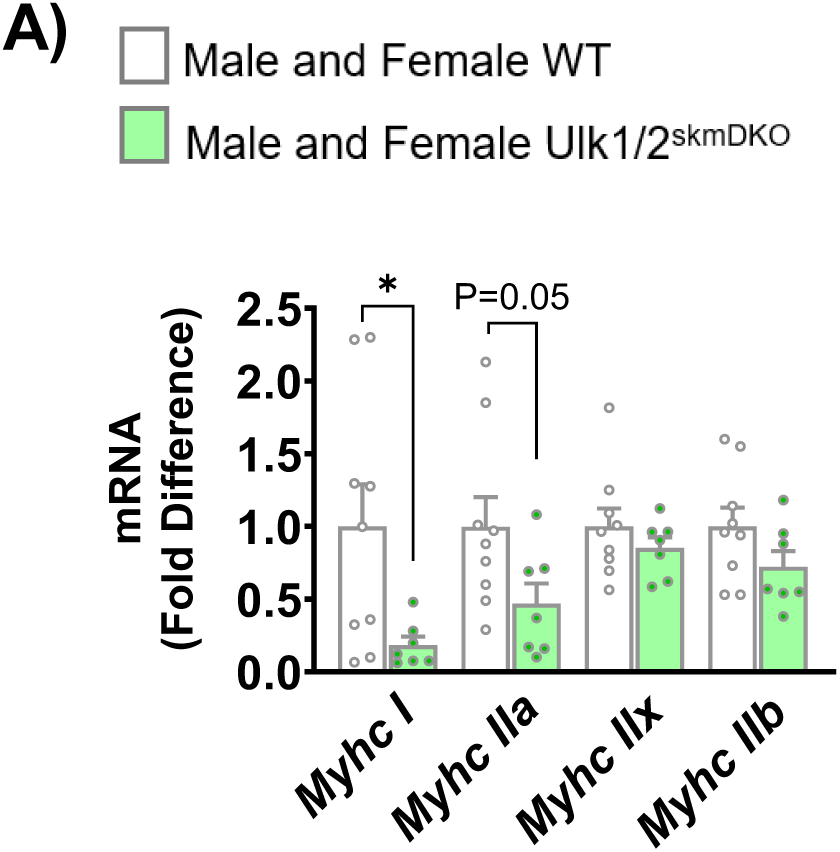
MyHC isoform mRNA expression. A) Relative mRNA expression of MyHC isoforms in tibialis anterior (TA) muscles from male and female mice (fold difference vs. WT, n = 7-9). Data represent means ± SEM. Statistical comparisons were performed using unpaired two-tailed *t*-tests. \**P* < 0.05.

**Supplementary figure 3.**
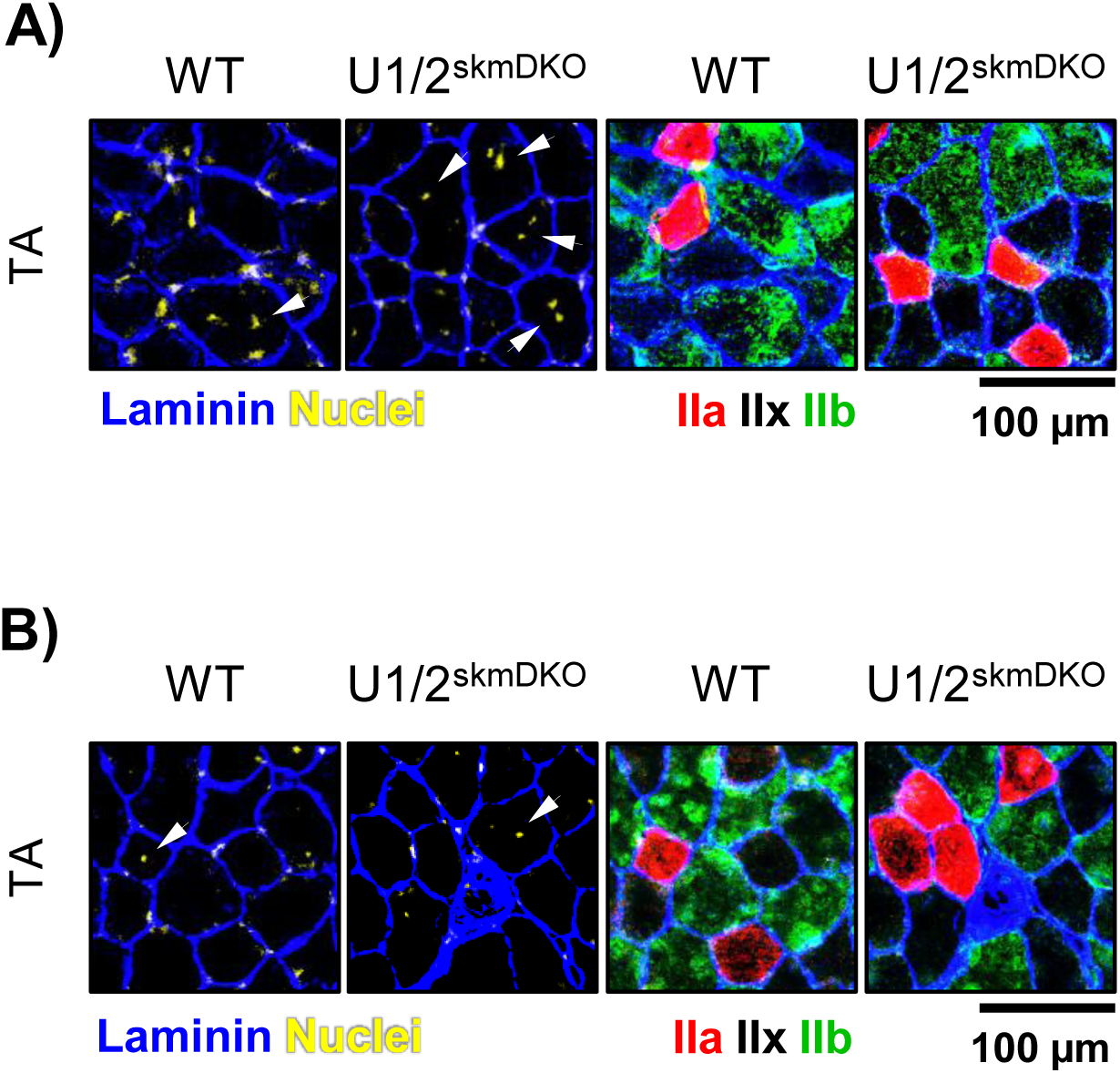
Centrally nucleated myofibers types in tibialis anterior muscle. A-B) Representative immunofluorescence images showing centrally nucleated (CN) myofibers, indicated by white arrowheads, in tibialis anterior (TA) muscles from male (A) and female (B) WT and skmDKO mice denoting laminin, nuclei and MyHC isoforms (Type IIa, IIx, IIb).

**Supplementary figure 4.**
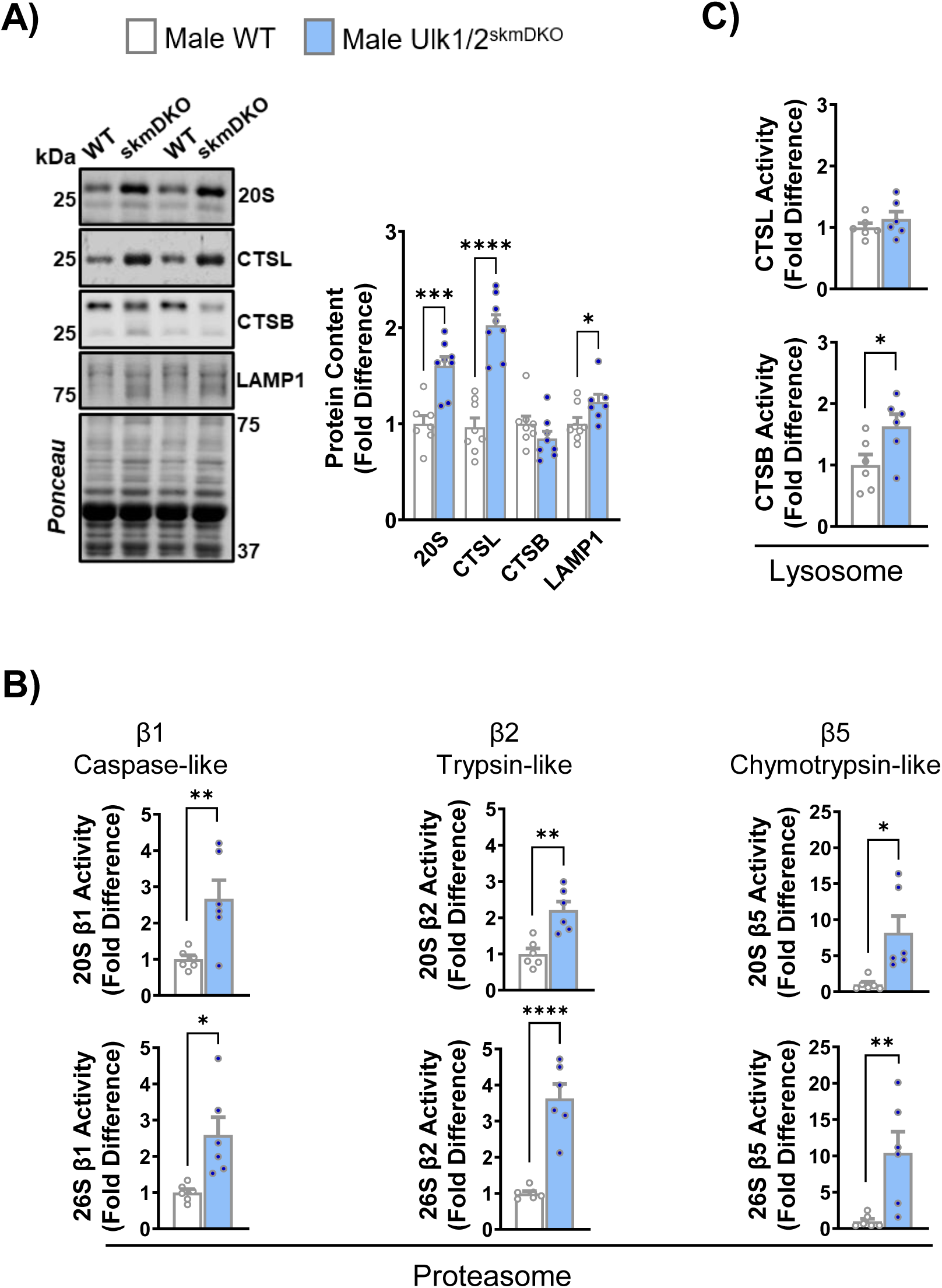
Proteasomal and lysosomal adaptations to muscle *Ulk1/2* loss. A) Representative immunoblots and quantification of 20S, CTSL, CTSB, and LAMP1 in tibialis anterior (TA) muscles (fold difference vs. WT, n = 7-8). B) Proteolytic activity β1 (caspase-like), β2 (trypsin-like), and β5 (chymotrypsin-like) subunits of the ATP-independent (20S) and ATP-dependent (26S) proteasome in gastrocnemius (GA) muscles (fold difference vs. WT, n = 6). C) CTSL and CTSB proteolytic activity in GA muscle (fold difference vs. WT, n = 6). Data represent means ± SEM. Statistical comparisons were performed using unpaired two-tailed *t*-tests. \**P* < 0.05, \*\**P* < 0.01, \*\*\**P* < 0.001, \*\*\*\**P* < 0.0001.

